# Targeting metabolic fluxes reverts metastatic transitions in ovarian cancer

**DOI:** 10.1101/2023.05.02.538518

**Authors:** Garhima Arora, Jimpi Langthasa, Mallar Banerjee, Ramray Bhat, Samrat Chatterjee

## Abstract

Spheroids formation during epithelial ovarian cancer progression correlates with peritoneal organ colonization, disease recurrence, and poor prognosis. Although cancer progression has been demonstrated to be associated with and driven by metabolic changes within transformed cells, possible associations between metabolic dynamics and metastatic morphological transitions remain unexplored. To address this problem, we performed quantitative proteomics to identify protein signatures associated with three distinct morphologies (2D monolayers and two geometrically individual three-dimensional spheroidal states) of the high-grade serous ovarian cancer line OVCAR-3. Integrating the protein states onto genome-scale metabolic models allowed us to construct context-specific metabolic models for each morphological stage of the OVCAR-3 cell line and systematically evaluate their metabolic functionalities. We obtained disease-driving metabolic reaction modules using these models and elucidated gene knockout strategies to reduce metabolic alterations associated with disease progression. We explored the DrugBank database to mine pharmacological agents and evaluated the effect of drugs in impairing cancer progression. Finally, we experimentally validated our predictions by confirming the ability of one of our predicted drugs: the neuraminidase inhibitor oseltamivir, to disrupt the metastatic spheroidal morphologies without any cytotoxic effect on untransformed stromal mesothelial monolayers.

## 1 Introduction

Epithelial ovarian cancer is one of the most common gynecological malignancies worldwide, with a high mortality rate and poor prognosis [1]. Global estimates of ovarian cancer reveal an incidence of three lakh new cases each year with an overall share of 1.6% of all cancers and an increased mortality rate [1, 2]. Delayed onset of clinical symptoms results in more than 70% of the cases left undiagnosed until cancer approaches an advanced stage with distant metastases. Auxiliary to this, the advanced stages of ovarian cancer culminate into a chemo-refractory form typically associated with poor overall survival [3, 4]. The metastasis of ovarian cancer begins with the detachment of transformed cells from the surface of the primary tumor due to decreased cell adhesion, and accumulation within the peritoneal cavity [5]. Changes in cell-cell adhesion follow a multi-step process of cancer progression, permitting epithelial cancer cells to migrate, remodel the extracellular matrix (ECM), and form clusters commonly known as spheroids. These spheroids contain cancer-associated fibroblasts and activated mesothelial cells, contributing to the development of the ascitic micro-environment [6]. The existence of spheroids within the ascitic fluid highly correlates with the recurrence of the disease and resistance to the first-line platinum-based chemotherapeutic drugs [7].

Alterations occurring in tumor cell mechanics require metabolic rewiring along this process to satisfy cancer cell’s energetic needs, which is one of the hallmarks of epithelial cancers. Studies with 13C-glucose reveal that the spheroid-forming cells with cancer stem-like properties (aggressive tumor growth, higher invasiveness, enhanced chemotherapy resistance) derive glucose predominantly through a combination of metabolic pathways to fulfill high anabolic demands [8]. The amino acids, namely, glutamine, glutamate, serine, and aspartate, which are essential for carrying out TCA cycle reactions, were significantly increased in spheroid-forming ovarian cancer cells compared to cancer cells cultured in adherent plates [9]. The mounting body of evidence recognized reprogramming of lipid metabolism as a hallmark of enhanced tumor metastasis and aggressiveness in ovarian cancer [10, 11]. Taken together, these represent significant findings demonstrating the connection between cell morphology and tumor progression with altered cell metabolism. This synergy unbolts the possibility of simultaneously targeting both the above aspects, thus halting disease progression more effectively.

Studies concerning metabolic alterations have used metabolic models to investigate different metabolic states of cells under normal and cancerous conditions and advance our ability to identify potential drug targets and biomarkers [12, 13, 14]. Pan-cancer analysis done by Gatto et al. [15], involving 13 different cancers, including ovarian, shows the general metabolic features separating normal and cancerous phenotypes. The study builds an insightful metabolism-driven narrative for cancer progression but does not explore tissue-specific cancer manipulations. Motamedian and coworkers have shown significant metabolic differences in the metabolism of cisplatin-sensitive and -resistant epithelial ovarian cancer cells and normal ovarian epithelia using a genome-scale metabolic model [16]. Evidently, few studies have been done on genome-scale metabolic models in ovarian cancer. In addition, to the best of our knowledge, there are no studies examining stage-or morphology-specific changes in metabolic dynamics of ovarian cancer.

In order to identify metabolic perturbations during the phenotypic transitions, we performed quantitative proteomics to delineate distinct protein signatures between monolayer cultures of an ovarian cancer cell line OVCAR-3 and its two distinct spheroidal morphologies (see Data File 1). Our objective was to predict the transition-specific perturbation modules responsible for disease progression and identify regulatory points that might revert the disease. We reconstructed context-specific metabolic models for OVCAR-3 samples of three morphological stages using Recon3D and examined the metabolic modulation associated with disease progression. We then explored the knockout strategies in these metabolic networks to reduce tumor-specific metabolic alterations during ovarian cancer progression. We investigated the DrugBank database to extract drugs against enzymes catalyzing the metabolic reactions and evaluated their effect in impairing ovarian cancer progression using the metabolic models of each spheroidal morphology. Finally, we experimentally validated our model predictions on the pharmacological regulation of metastatic spheroidal morphologies.

## 2 Material & Methods

### 2.1 Data acquisition

The ovarian cancer cell line OVCAR-3 (obtained from ATCC, Virginia, US) was used for this study. Cells were maintained and cultured in suspension following the protocol used in our previous study [17]. Quantitative mass spectrometry was performed on three independent biological replicates of each of the distinct phenotypic morphologies (see Supplementary Information section: S1.1). The proteomics data obtained contains the expression of 3122 proteins across triplicates of the different morphologies of OVCAR-3. This data was further pre-processed to filter metabolic genes data in order to construct metabolic models (details given in Supplementary Information section: S1.2).

### 2.2 Constructing context-specific metabolic models

We built the context-specific metabolic models for each morphological stage using model extraction methods (MEMs) namely, iMAT, FASTCORE, INIT. The CBM sampling technique, GP sampler was used to produce a sequence of feasible solutions and the solution having a maximum correlation with the reaction’s expression array (see Supplementary Information section: S1.3.2) was regarded as the optimal flux solution. The detailed information from constructing context-specific metabolic models to evaluating optimal flux solution is provided in Supplementary Information section: S1.3 and for complete workflow see Fig. S1.

### 2.3 Metabolic functionality score’s significance

We assessed the significance of the metabolic functionality score evaluated for each context-specific model. We computed the metabolic functionality score of 1000 random models of the same size as the queried model under investigation. These models were generated by randomly removing reactions from the parent model (Recon3D) while ensuring the model’s consistency. Then we defined the p-value as follows:

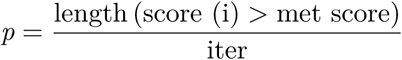

where, score is the metabolic functionality score for the i^*th*^ randomly extracted model from Recon3D, met score is the metabolic functionality score of the queried model, and iter is the number of randomization (i.e., 1000 in our case).

### 2.4 Reaction modulation approach

Reaction modulation aims to find modules of reactions whose perturbation is associated with disease progression. Reactions perturbed during morphological transitions were considered for single reaction modulation, where each reaction was modulated individually, and the correlation between the flux states before and after modulation was evaluated against each perturbed reaction. A separate model was built corresponding to each perturbed reaction where that particular reaction was modulated, i.e., the upper and lower bounds of that reaction were constrained using corresponding flux values of that reaction in the initial stage model of the transition. The perturbed reactions were sorted based on their correlation coefficient obtained from the single reaction modulation technique. We used models obtained from the single reaction modulation and started modulating the perturbed reactions sequentially based on their correlation values found during the single modulation technique. The correlation coefficient was evaluated on the modulation of each new reaction in the model, and the modulation was only considered significant if it increased the correlation by 0.001. Otherwise, the modulation of that particular reaction was not added to the model. Following this way, the iteration goes up to the length of the total number of perturbed reactions. The resultant model and the corresponding correlation coefficient were again put to another level of sequential reaction modulation in the same manner. This is defined as different levels of reaction modulation. So iteratively, in each level, if the sequential modulation of a reaction results in a significant increase in correlation coefficient, it was added to the model, and the model was sent to the next level until the correlation between consecutive levels becomes less than 0.01.

### 2.5 Filtering non-essential genes

The Genotype-Tissue Expression (GTEx) database was used to extract RNA sequence data of 166 normal ovarian samples [18, 19]. The data were quantile normalized, which ensured that the distributions were identical across samples. We then used the E-flux approach to integrate data into Recon3D, generating 166 metabolic models, one for each normal ovarian sample. The bounds of reactions catalyzed by the queried gene were set to zero in order to see its knockout effect on the growth rate of normal ovarian models. The Flux balance analysis (FBA) was then performed using Cobratoolbox’s function ‘Optimizecbmodel’ to determine the growth rate [20]. This procedure was repeated for each gene in each of the normal ovarian models described above. The ratio of growth rates before and after gene deletion was then calculated for each gene in all normal ovarian models. Genes with a growth rate ratio of 1 in at least one-third of the total normal ovarian models were classified as non-essential since knocking them out did not influence the growth rate of normal ovarian models.

### 2.6 Approximating minimal percentage inhibition by drug targets

The aim is to determine the minimum percentage inhibition or reduction in the flux bound of reactions catalyzed by metabolic targets of the drug that can produce a maximum effect, i.e., a high correlation coefficient. We varied the percentage inhibition of the flux bounds from 10% (partial inhibition) to 100% (complete inhibition). For both transitions, the correlation coefficient between the flux values of the perturbed reaction in the drug infused metabolic model and the initial stage model was evaluated. The extent of positive correlation indicates the drug’s potential in reverting the transition.

### 2.7 Cell culture and reagents

OVCAR-3 cells were maintained in RPMI-1640 (Roswell Park Memorial Institute) medium (HiMedia, AL162A) along with 10% FBS (Gibco, 10270). The non-cancerous mesothelial cells, MeT-5A were cultured in complete Medium 199 supplemented with 10% FBS, 5 *μ*g/ml insulin, 0.5 *μ*g/ml hydrocortisone, 2.6 ng/ml sodium selenite, 27 pg/ml *β*-estradiol, 10 *μ*g/ml transferrin, 10 ng/ml hEGF and 20 mM HEPES buffer. All the cells were maintained in a 5% CO_2_, 37°C temperature humidified incubator. Oseltamivir phosphate (SML1606) was the drug used in our study.

### 2.8 Spheroid culture

Spheroids were cultured in 3% polyHEMA-coated (Sigma, P3932) dishes in RPMI-1640 defined medium supplemented with 250 ng/ml insulin (Sigma-Aldrich, I6634), 0.5 *μ*g/ml hydrocortisone (Sigma-Aldrich, H0888), 2.6 ng/ml sodium selenite (Sigma-Aldrich, S5261), 10 *μ*g/ml transferrin (Sigma-Aldrich, T3309), 27.3 pg/ml estradiol (Sigma-Aldrich, E2758) and 5 *μ*g/ml prolactin (Sigma-Aldrich L6520). These were maintained in culture for 1 to 7 days. Spheroids were visualized using the Olympus IX73 fluorescence microscope.

### 2.9 Cell staining and imaging

Spheroids were first pelleted in a sterile 15-ml tube. Cells were then fixed using 3.7% formaldehyde (24005; Thermo Fisher Scientific) at 4°C for 30 min. After fixation, cells were washed and resuspended in PBS. Following this, 20 *μ*l of spheroid suspension was dried in eight well chambered cover glass at 37°C for 30 mins. Permeabilization was achieved using 0.5% Triton X-100 in PBS (PBST) (MB031; HiMedia) for 2h at RT. Alexa Fluor^*T M*^ 633-conjugated phalloidin (Thermo Fisher Scientific, A22284) was added to cells at 1:500 dilution in 0.1% PBST and incubated overnight at 4°C. Cells were washed thrice with 1 × PBS for 5 mins, counterstained with 1 *μ*g/mL DAPI (D1306; Thermo Fisher Scientific).

To determine live and dead cells in the culture, cells were stained with 0.5 mg/ml Calcein AM (C1430, Invitrogen) for 15 mins. Cells were washed with PBS, counterstained with Propidium Iodide (TC252, Himedia) for 5 mins, and imaged under the Olympus IX73 fluorescence microscope.

## 3 Results

### 3.1 Generation of context-specific metabolic models for each morphological stage

Differentially expressed proteins between the different morphological stages of the OVCAR-3 cell line were identified using the acquired protein expression data (see Material & Methods: 2.1). Substantial involvement of enzymatic genes such as GLS, DBI during transition 1 and ILK, MAP2K1 during transition 2 was observed (for details, see Supplementary Information section: S2.1 and Fig. S2). The pathway enrichment analysis of these altered proteins uncovered the significant involvement of metabolic pathways in disease progression (see Data File 2). To acquire a better understanding of how the morphological changes during tumor progression are associated with metabolic reprogramming, we used a generic human metabolic model Recon3D [21]. Choosing an efficient model extraction method (MEM) to create context-specific models has always been challenging because it has a considerable impact on the size and functionality of the context-specific models [22]. As we know, cells perform distinct metabolic functions under different physiological conditions; hence pre-defining a specific metabolic objective of a cell for all conditions is not appropriate. Moreover, previous experimental observation also suggested that during the transition from moruloid spheroids to blastuloid spheroids, the phenotypes did not exhibit any significant difference in cell proliferation [17]. Therefore, to build metabolic models for each morphological stage of the OVCAR-3 cell line, we have used three different MEMs that are independent of the objective function and each with two different thresholds (see Material & Methods: 2.2). The number of reactions and metabolites present in all metabolic models built for each morphological stage is given in Data File 3.

The central premise of these MEMs is to preferentially extract reactions corresponding to highly expressed genes. However, the cellular metabolic functionalities are not always preserved after model extraction. The study by Opdam et al. [22] provided 56 metabolites based on the biomass function that are essential for cancer cell growth. These metabolites include the synthesis of non-secreted metabolites, e.g., glutathione, ATP, and carnitine. Following this study, we evaluated the metabolic functionality score for all the afore-built context-specific models (see Fig. 1a). Furthermore, a multivariate PCA analysis of the steady state reaction’s flux values (see Supplementary Information section: S1.4) in all the models revealed their dependencies on the choice of MEM and threshold (see Fig. 1b, 1c). It was observed that the models built using the iMAT method had a high metabolic functionality score and were less sensitive to the threshold selection. For all three morphological stages of the OVCAR-3 cell line, iMAT models with threshold 1 (iMAT-Th1) were chosen to gain further deep insights into the metabolic liabilities. The flux values of sink reactions corresponding to 56 metabolites based on biomass function in all three morphological stage-specific iMAT-Th1 models are illustrated in Fig. 1d. During the phenotypic transition from monolayer to spheroidal morphologies, the flux values of the sink reaction associated with the metabolite cholesterol increased significantly. The flux values of sink reaction corresponding to asparagine, threonine, leucine, and valine, on the other hand, decreased. The metabolic profiling of iMAT-Th1 models revealed the perturbation of 1226 and 1252 reactions during transition 1 and transition 2, respectively (see Supplementary Information section: S2.2). The metabolic pathway enrichment analysis was carried out using the Recon3D database, and the findings demonstrated a substantial role of lipid, carbohydrate, and protein/amino-acid metabolism in the disease progression (see Fig. S3).

**Figure 1:**
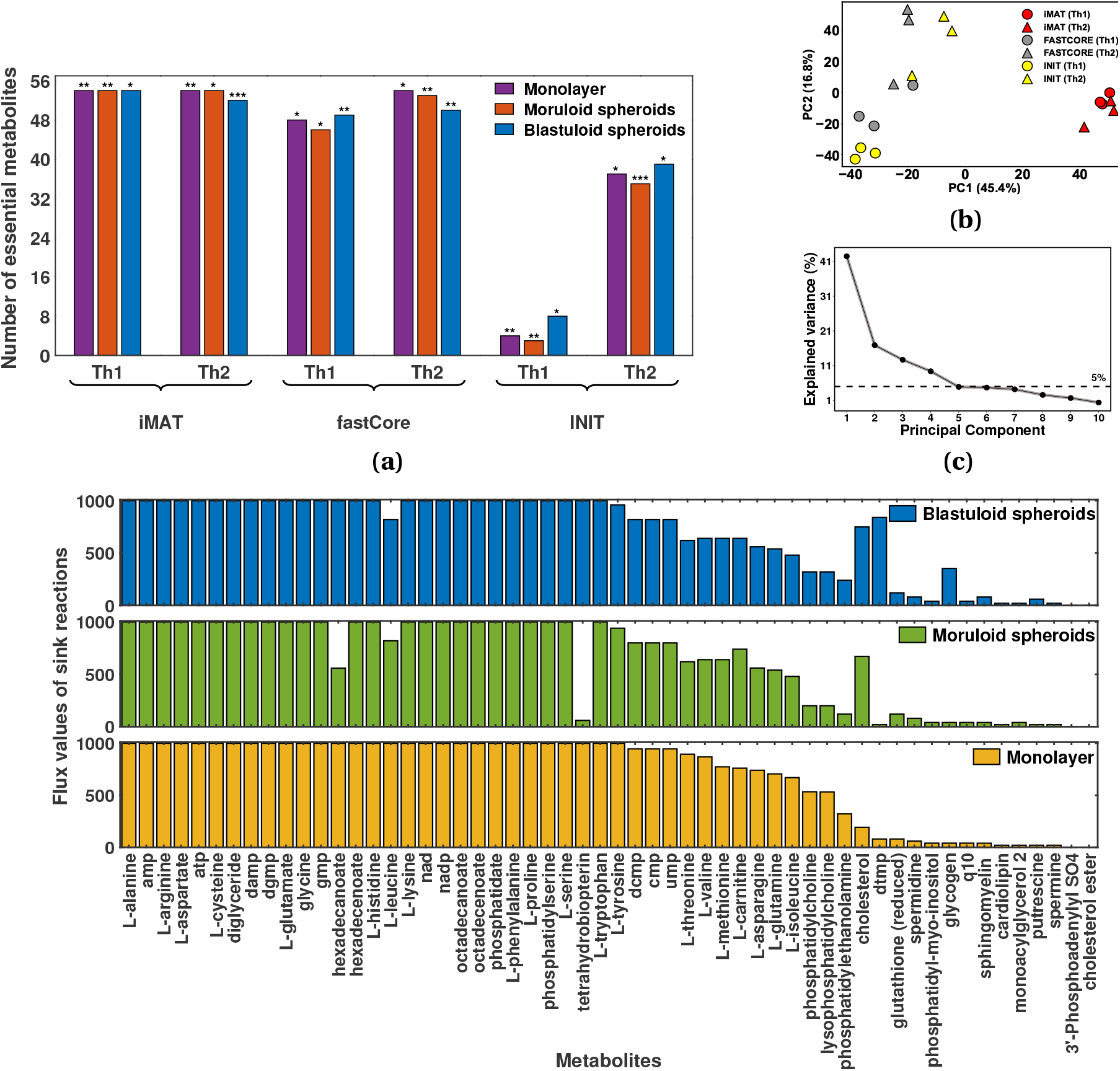
Systematic evaluation of different Model Extraction Methods (MEMs): (a). Bar graph shows significant metabolic functionality scores, i.e., the number of essential metabolic functions for cancer growth having non-zero flux values, corresponding to each model. The significance of the metabolic functionality score (see Material & Methods: 2.3) for each model is marked at the top of each bar (*p*<*0.05, **p*<*0.01, ***p*<*0.001). (b). PCA plot represents the clustering of metabolic models for different morphological stages of the OVCAR-3 cell line based on different MEMs and threshold values. The red dots in the PCA plot represent models built using the iMAT technique, whereas the grey and yellow dots represent models built using FASTCORE and INIT, respectively. (c). The line plot depicts the percentage variance explained in each principal component. (d). Bar graphs showing flux profiles of sink reaction corresponding to each metabolite essential for cancer growth in iMAT-Th1 models of each morphological stage.

### 3.2 Identification of core reaction modules associated with disease progression

Metabolic alterations drive OVCAR-3 cancer cells to gain a metastatic phenotype and form spheroids in the peritoneal cavity. Perturbations directing the transition of morphological stages of the OVCAR-3 cell line were obtained in previous section. A low correlation of 0.0284 and 0.0352 ascertained the significant shift in the metabolic flux profiles during the transition 1 and transition 2, respectively. Both common and transition-specific reactions were large in number, and targeting such large perturbations is biologically infeasible. Therefore, we defined a novel approach for finding an optimal set of metabolic reactions whose perturbation is responsible for the transition of the morphological stages of the OVCAR-3 cell line. This technique was aimed at finding small modules of reactions whose modulation increases the correlation between the flux profiles of perturbed reactions in the modulated and initial stage model of the transition (see Fig. 2a). In single reaction modulation, the mean correlation between the perturbed flux profiles during transition 1 and transition 2 were 0.0767 and 0.0896, respectively, which is not a very significant increase. This reiterated the fact that the disease alters the cellular metabolism at multiple sites; hence, simultaneous modulation of multiple reactions should be an optimal approach. So, we performed multiple reaction modulation (see Material & Methods: 2.4), and the maximum correlation achieved for both transitions were 0.4744 and 0.5151, respectively. The number of modulated reactions and the associated correlation coefficient obtained after modulation were found to have no relationship (see Fig. 2b, 2c).

**Figure 2:**
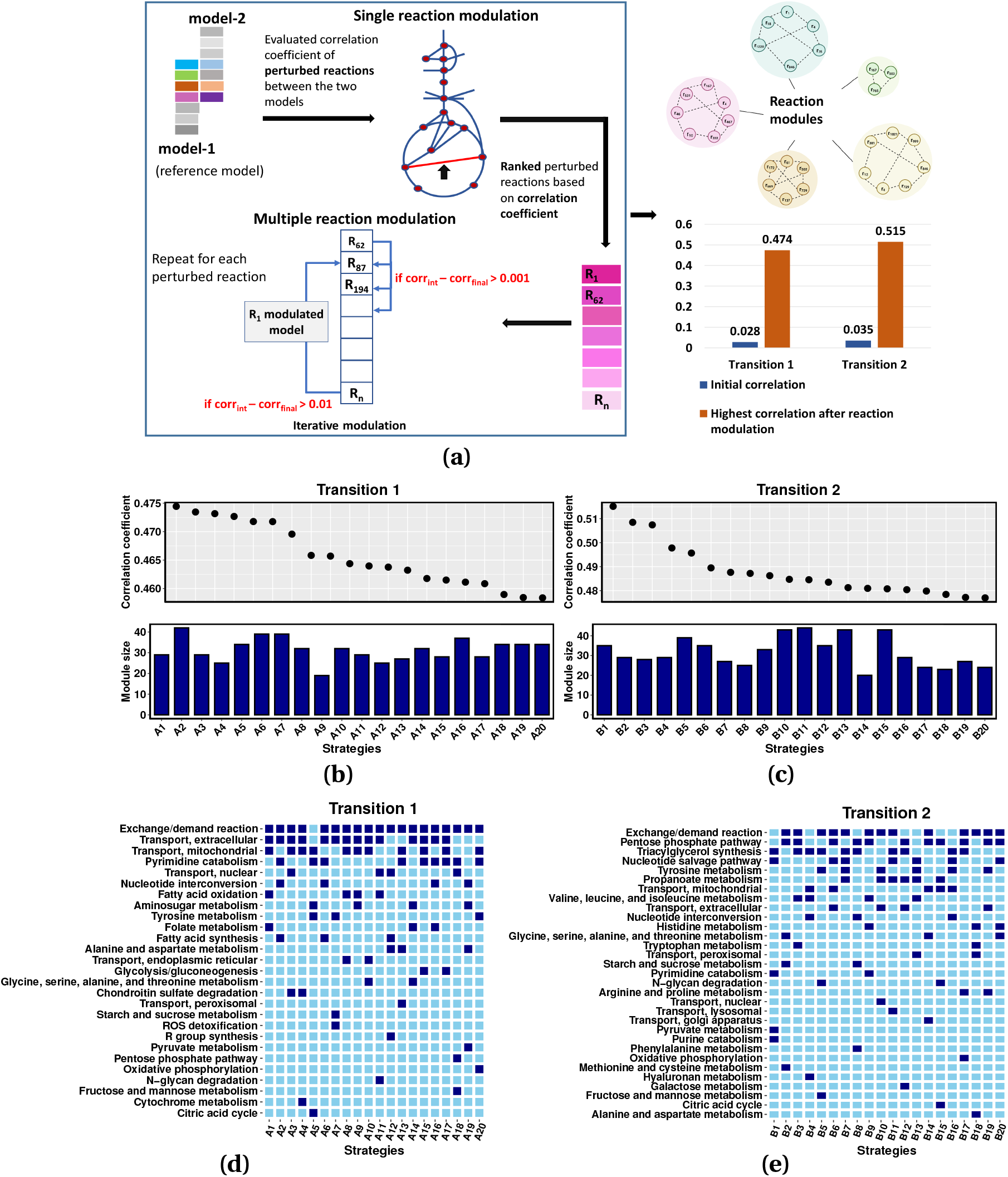
Grouping metabolic perturbed reactions based on their influence on morphological transitions: (a). Schematic diagram summarizes the reaction modulation process followed during the search for perturbed reaction modules responsible for the disease transition. (b),(c). The figure shows the correlation coefficient obtained corresponding to the top 20 reaction modulation strategies together with the number of reactions modulated in each strategy. (d),(e). The plot shows the pathways involved in the top 20 modulation strategies (Ai and Bi, i=1,2,..,20) for both transitions. The involvement of a pathway in a particular strategy is represented by dark blue color, whereas light blue color represents its absence.

The reactions present in top modulation strategies (see Data File 4) of both the transitions were involved in exchange/demand reaction pathway, and a few transport pathways such as transport extracellular, transport mitochondrial (see Fig. 2d, 2e). Carbohydrate related pathways such as citric acid cycle (TCA), pentose phosphate, oxidative phosphorylation, pyruvate metabolism, and pathways involved in amino acid metabolism such as tyrosine, alanine & aspartate metabolism were involved in top modulation strategies of both the transitions. In addition, when comparing transition 2 to transition 1, reactions involved in the pentose phosphate pathway were found in many of the top modulation strategies. Only transition 1’s top modulation strategies showed evidence of glycolysis/gluconeogenesis, fatty acid oxidation, and synthesis. The involvement of the tri-acyl glycerol synthesis pathway (TAG) was observed in 10 of the top 20 modulation strategies for transition 2. Arginine and proline pathway involved in amino acid metabolism was also involved in transition 2’s top modulation strategies.

### 3.3 Predicting novel drug targets in reversing the metabolic alterations during OVCAR-3 morphological transitions

Alterations in the flux values of metabolic reactions were observed during the transition from the initial stage to the later stages of OVCAR-3. Targeting genes catalyzing the perturbed reactions can be a potential strategy for reversing the metabolic alterations that occur during morphological transitions. For this, we filtered out a total of 651 and 626 genes catalyzing reactions that were perturbed during transition 1 and transition 2, respectively. We sifted 495 and 462 non-essential genes (see Material & Methods: 2.5) to perform single gene knockouts in the iMAT-Th1 models of moruloid and blastuloid spheroids, respectively. The metabolic flux profiles after single gene knockout were then examined and compared to the flux profiles of the transition’s initial stage model. After executing single gene knockout using genes catalyzing reactions altered during transitions 1 and 2, the highest correlation coefficients obtained were 0.31447 and 0.33119, respectively (see Fig. 3a, 3b).

**Figure 3:**
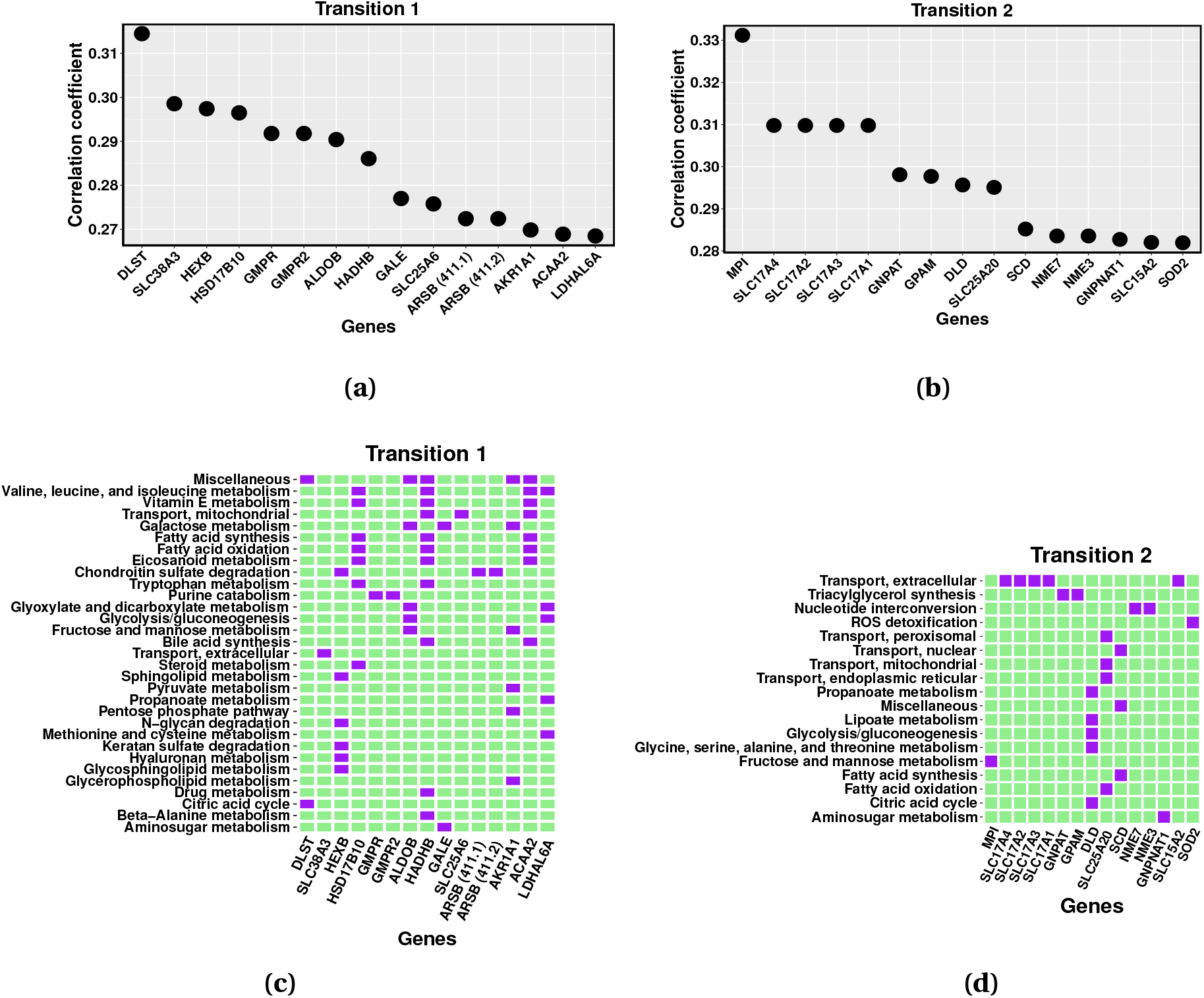
Identification and *in silico* knockout of targets identified through correlation analysis and their catalyzed pathways: (a),(b). The figure shows the top 15 gene knockout targets based on the correlation coefficient. (c),(d). Pathways enriched by reactions catalyzed by the top 15 gene knockout targets are depicted in the figure. The gene’s involvement in the corresponding pathway is represented by purple, whereas its absence is represented by green.

Silencing of genes involved in carbohydrate metabolism pathways, namely, DLST, GALE showed a significantly high correlation, i.e., were able to rectify the metabolic profile altered during transition 1. Multi-pathway targeting genes HSD17B10, HADHB, and ACAA2, were among the top 5 gene knockout results of transition 1 (see Fig. 3c). Silencing of many solute carrier (SLC) groups of membrane transport proteins, such as SLC17A4, SLC17A2, SLC25A1, and others (see Data File 5), were able to reduce the metabolic perturbations occurred during transition 2. Interestingly, knocking down of genes GNPAT and GPAM implicated in the tri-acyl glycerol synthesis pathway (lipid metabolism) were among the top gene knockout results of transition 2 (see Fig. 3d). This follows our finding in the previous section that the tri-acyl glycerol pathway undergoes a significant amount of perturbation, as its reactions were prominent among the top reaction modulation strategies of transition 2. We also observed that a single gene knockout of ALDOB, ADH1B, NEU1, and NT5E minimized the metabolic changes during both transitions.

Proteins essential to cancer survivability are of great interest in the search for new therapeutic targets [23, 24]. To investigate the effect of targeting such essential protein encoding genes in ovarian cancer, we retrieved 36 metabolic genes from the study by Kanhaiya et al. [25]. These genes were subjected to single gene knockout in our iMAT-Th1 models of moruloid and blastuloid spheroids of OVCAR-3. We observed an increase in correlation between the flux profiles of perturbed reactions in the acquired model following gene knockout and the initial model of the transition, i.e., log_2_ fold-change in correlation was greater than 2. Ensuing gene essentiality results, we assigned a targetability score to the top 20 reaction modulation strategies of both transitions. The percentage of the non-essential genes involved in each modulation strategy was defined as the targetability score (see Tables S1, S2).

### 3.4 Investigating the effect of drugs in reversing back the disease progression

Drug repurposing, also known as re-profiling, is instrumental in exploring new therapeutic uses for approved or investigatory drugs for a new medical indication. To perform in-silico drug repurposing in the models of moruloid and blastuloid spheroids, we extracted the information of 13,000 drugs and their corresponding targets from the DrugBank database [26]. We were able to map targets of 121 and 125 drugs (inhibitors) to the metabolic reactions catalyzed by them, respectively, in models of moruloid and blastuloid spheroids (see Fig. 4a). The effect of a drug (inhibitor) in the models was evaluated by reducing the flux bounds of reactions catalyzed by the metabolic targets of the drug. It was observed that the mean correlation coefficient obtained from the drug infused models increased up to 50% inhibition, and a further increase in percentage inhibition has little or no effect on the flux profiles (see Fig. 4b, 4c). Therefore, 50% reaction bound inhibition was chosen to achieve the optimum efficacy during drug repurposing (see Material & Methods: 2.6). For each drug, the correlation coefficient between the flux values of perturbed reactions in drug infused model and the initial stage model of a transition was evaluated (see Data File 6).

**Figure 4:**
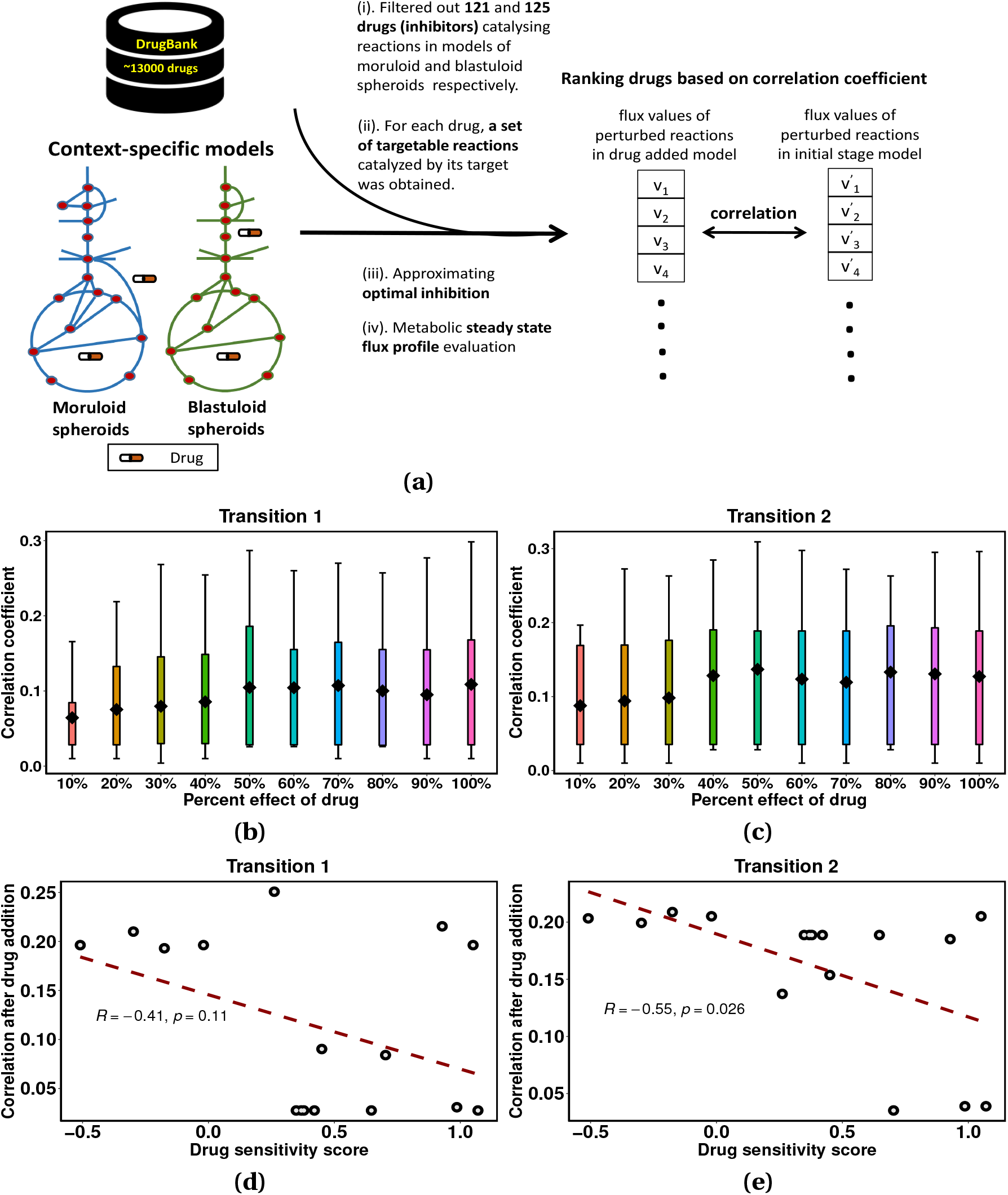
Estimation of the effect of repurposable drugs in altering the metabolic state of ovarian cancer cells using the metabolic models: (a). The workflow for evaluating the effect of drug response in the metabolic models. (b),(c). The figure shows the percentage effect of the drug over the model readouts after adding the drug. (d),(e). The regression plot shows an inverse relationship between the model readout, i.e., the correlation coefficient corresponding to 16 drugs and the drug’s sensitivity score obtained from GDSE. A positive score implies low sensitivity of the OVCAR-3 cell line, whereas a negative score implies a high sensitivity of the OVCAR-3 cell line towards the drug administration.

We filtered out drugs having targets overlapped with the gene list utilized for gene knockout and having log_2_ fold-change in correlation greater than 2 (see Table 1, 2). Among the drug repurposing results, single gene targeting drugs, namely disulfiram, ethoxzolamide, gadopentetate dimeglumine, and tolcapone, were specific to transition 1, whereas amphetamine, cefdinir, cerulenin were specific to transition 2. The single gene targeting drugs canagliflozin, moexipril, oseltamivir, myo-inositol, glyburide, gemcitabine, pentoxifylline, and salicylic acid target the genes involved in metabolic alterations occurring in both the disease transitions. Few multi-target drugs, such as benzthiazide, clodronate, fomepizole, hexachlorophene, methyclothiazide, and troglitazone, with all their targets involved in the reactions perturbed during transition 1. Clodronate and fomepizole, on the other hand, were the only two multi-target drugs, having all of their targets present in the metabolic perturbation developed during transition 2.

**Table 1:** Drug candidates targeting metabolic perturbations during transition 1.

| Drugs | Target<br>(Gene ID) | Target<br>(Gene Name) |
| --- | --- | --- |
| Glyburide | 19.1 | ABCA1 |
| Gemcitabine | 51727.1 | CMPK1 |
| Canagliflozin | 6523.1 | SLC5A1 |
| Disulfiram | 217.1 | ALDH2 |
| Tolcapone | 1312.1 | COMT |
| Moexipril | 59272.1 | ACE2 |
| Ethoxzolamide | 766.1 | CA7 |
| Gadopentetate dimeglumine | 5226.1 | PGD |
| Oseltamivir | 4758.1 | NEU1 |
| Myo-Inositol | 10423.1 | CDIPT |
| Pentoxifylline | 4907.1 | NT5E |
| Salicylic acid | 1645.1 | AKR1C1 |
| Benzthiazide | 771.1, 768.1 | CA12, CA9 |
| Clodronate | 291.1, 292.1, 293.1 | SLC25A4, SLC25A5, SLC25A6 |
| Fomepizole | 124.1, 125.1, 126.1, 847.1 | ADH1A, ADH1B, ADH1C, CAT |
| Hexachlorophene | 2746.1, 6392.1 | GLUD1, SDHD |
| Methyclothiazide | 759.1, 760.1, 762.1 | CA1, CA2, CA4 |
| Troglitazone | 2182.1, 2030.1 | ACSL4, SLC29A1 |

**Table 2:** Drug candidates targeting metabolic perturbations during transition 2.

| Drugs | Target<br>(Gene ID) | Target<br>(Gene Name) |
| --- | --- | --- |
| Canagliflozin | 6523.1 | SLC5A1 |
| Cerulenin | 2194.1 | FASN |
| Cefdinir | 4353.1 | MPO |
| Moexipril | 59272.1 | ACE2 |
| Oseltamivir | 4758.1 | NEU1 |
| Myo-Inositol | 10423.1 | CDIPT |
| Amphetamine | 4128.1 | MAOA |
| Glyburide | 19.1 | ABCA1 |
| Gemcitabine | 51727.1 | CMPK1 |
| Pentoxifylline | 4907.1 | NT5E |
| Salicylic acid | 1645.1 | AKR1C1 |
| Clodronate | 291.1, 292.1, 293.1 | SLC25A4, SLC25A5, SLC25A6 |
| Fomepizole | 124.1, 125.1, 126.1, 847.1 | ADH1A, ADH1B, ADH1C, CAT |

We compared the predicted drug response evaluated through our models with the drug sensitivity scores (IC50 z-score) obtained from Genomics of Drug Sensitivity in cancer (GDSE) for the OVCAR-3 cell line [27]. We obtained the sensitivity score for 16 drugs mapped with our drug list (see Tables S3, S4). The correlation coefficient corresponding to the mapped drugs and their sensitivity score showed an inverse relationship suggesting that the OVCAR-3 cell line will be more sensitive toward the drugs with a high correlation coefficient (see Fig. 4d, 4e). Single-target drugs common to both transitions was chosen (from Table 1, 2) and pathway enrichment analysis was carried out to gain mechanistic insights using the reactions catalyzed by the targets of these drugs. It was observed that the drug oseltamivir targeting neuraminidase was involved in highest number of pathways as compared to other drugs (see Data File 7). We also evaluated the overlap value for each pathway, which is calculated as the ratio of number of reactions targeted by the drug in that pathway by the total number of reactions in that pathway. Oseltamivir showed larger overlap values and higher pathway perturbation than the other drugs.

### 3.5 Experimental validation of drug repurposing predictions

To validate our prediction on the pharmacological regulation of both transition 1 and transition 2, we chose oseltamivir, which is present in both transitions with large overlap values and high pathway perturbation. It targets neuraminidase which has been demonstrated to contribute to the progression of various cancers [28, 29, 30]. Although the levels of Neuraminidase-1 have been correlated with ovarian cancer cell proliferation and invasion [31], an association between the ability of its enzymatic activity to regulate the sialic acid flux and the formation of spheroids, the principal mediators of ovarian cancer metastasis [32], as revealed through the global metabolic analysis of the spheroidal proteomics is as yet unknown. So, we chose this drug for further *in vitro* evaluation in ovarian cancer. Progressively increasing concentrations of oseltamivir (100 *μ*M - 5 mM) were added to suspended single OVCAR-3 cells, which were cultured for 24-hours. The concentration of oseltamivir was chosen based on pharmacological literature [33, 34, 35]. Control cells formed moruloid spheroids within this time period. On the other hand, moruloid spheroid formation (transition 1) was impaired in oseltamivir-added cultures upon treatment with concentrations of 250 *μ*M and upwards (morphology and cellularity assessed using staining for F-actin using phalloidin and for DNA using DAPI; 2 mM EGTA was used as a positive control as it manages to impair and disrupt spheroid formation see Fig. 5a and Fig. S4a for phase contrast images). Upon treatment of already-formed moruloid spheroids with progressively increasing concentrations of oseltamivir, disruption of transition 2, was observed from 250 *μ*M and upwards (morphology and cellularity assessed using staining for F-actin using phalloidin and for DNA using DAPI; 2 mM EGTA was used as a positive control as it manages to impair and disrupt spheroid formation see Fig. 5b and Fig. S4b for phase contrast images). We next probed whether the impairment of transition 1 and 2 was a result of dose-dependent cytotoxicity exerted by oseltamivir. When spheroids were treated similarly as in experiments of Fig. 5a, 5b, and stained instead for Calcein AM (to detect viability) and proidium iodide incorporation (to detect cell death), we found that although disrupted, neither impairment nor disintegration of spheroids were accompanied by cell death suggesting the drug exerted its effect through disrupting cellular interactions and not through cell death (Fig. 6a, 6b). We next asked if the spheroidogenesis-disrupting concentrations of oseltamivir have any effects on untransformed mesothelial cells that constitute the tissue micro-environment associated with ovarian cancer metastasis. We observed no cytotoxic effects exerted by 250 *μ*M and 500 *μ*M oseltamivir (as assessed through Calcein AM and propidium iodide treatment; Fig. 7). Our results, therefore, confirm the GSMM-based predictions that oseltamivir can be a potent disruptor of ovarian cancer spheroidogenesis and its heterogeneous transitions, without exerting any side effects on the non-cancerous cells similar to those that line the peritoneal cavity within which metastasis takes place.

**Figure 5:**
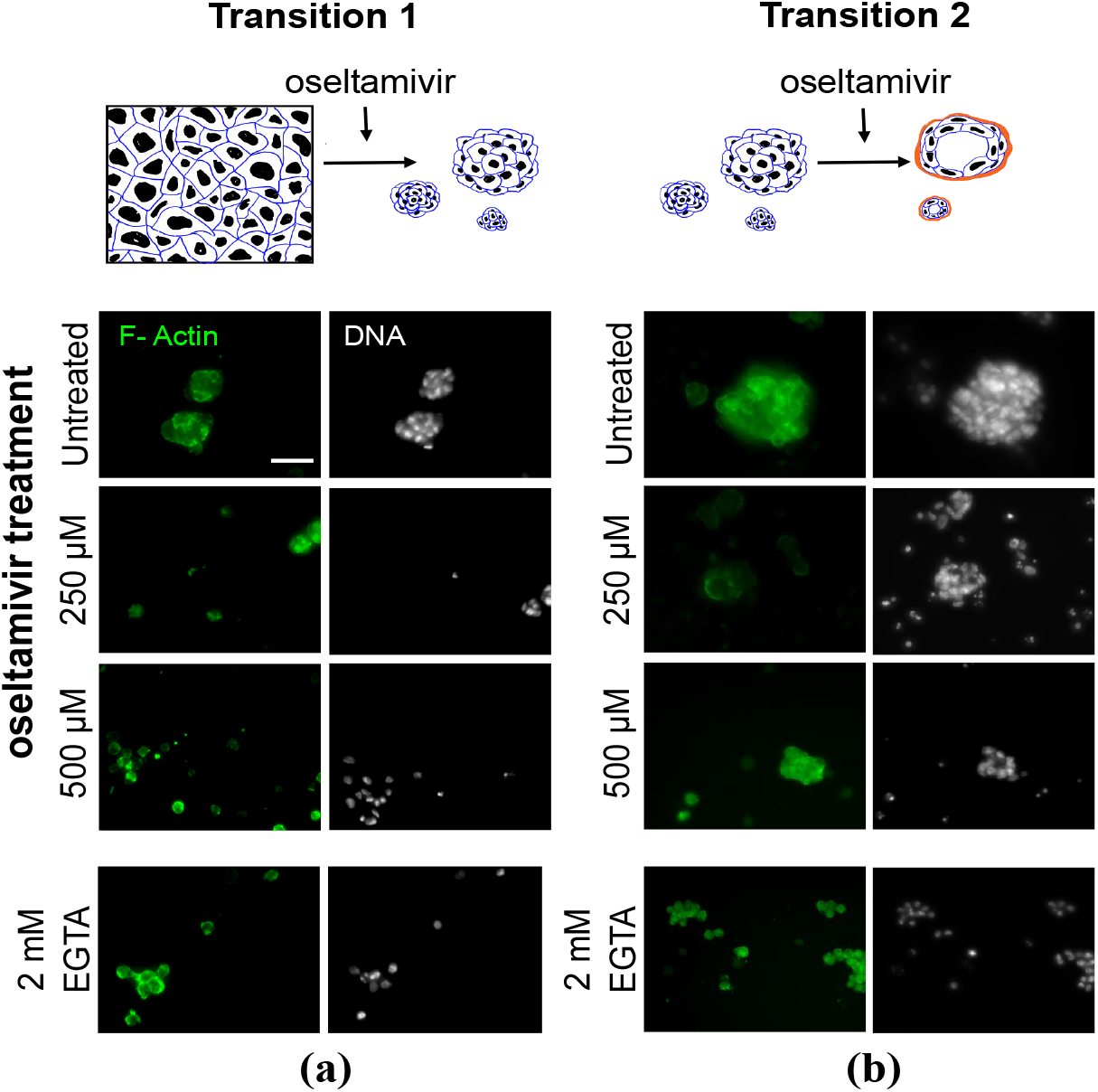
Oseltamivir treatment disrupts single cell-to-moruloid and moruloid-to-blastuloid transitions of ovarian cancer spheroids: (a). Laser confocal photomicrographs of ovarian cancer OVCAR-3 cells allowed to transit from single cells to moruloid spheroids (transition 1; top represents a cartoon representation of the same). Cells were untreated (top row of micrographs) or treated with 250 *μ*M oseltamivir (second from top), 500 *μ*M oseltamivir (third from top), and 2 mM EGTA (bottom row) and observed after 24 h of culture with staining for F-actin (green using phalloidin) and DNA (white using DAPI) (n = 3). (b). Laser confocal photomicrographs of ovarian cancer OVCAR-3 cells allowed to transit from moruloid to blastuloid spheroids (transition 2; top represents a cartoon representation of the same). Spheroids were untreated (top row of micrographs) or treated with 250 *μ*M oseltamivir (second from top), 500 *μ*M oseltamivir (third from top), and 2 mM EGTA (bottom row) and observed after 24-72 h of culture with staining for F-actin (green using phalloidin) and DNA (white using DAPI) (n = 3). Scale bar: 50 *μ*m.

**Figure 6:**
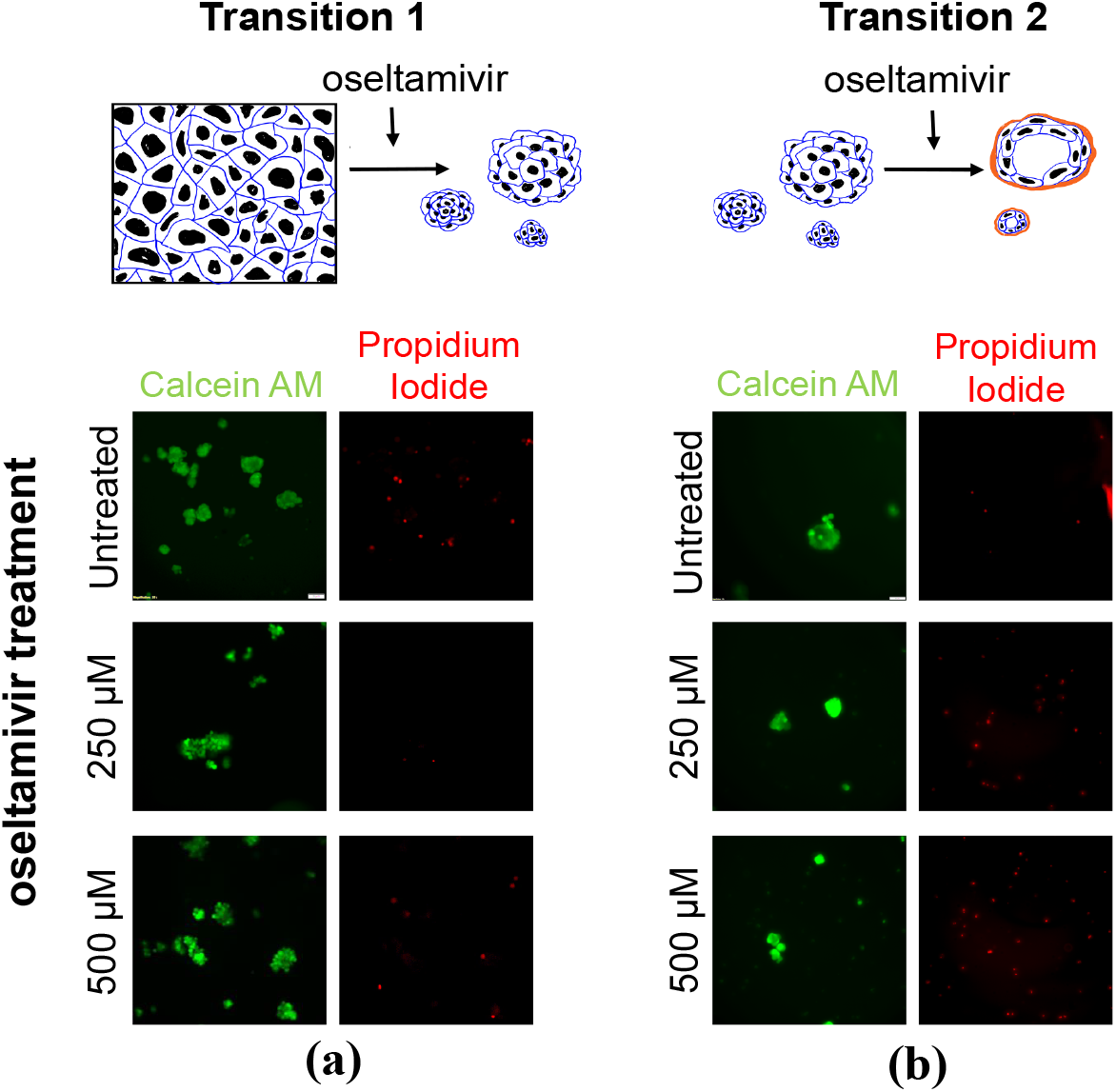
The inhibition of spheroidogenesis by oseltamivir is not accompanied by cell death: (a). Laser confocal photomicrographs of ovarian cancer OVCAR-3 cells allowed to transit from single cells to moruloid spheroids (transition 1; top represents a cartoon representation of the same). Cells were untreated (top row of micrographs) or treated with 250 *μ*M oseltamivir (second from top), 500 *μ*M oseltamivir (third from top), and 2 mM EGTA (bottom row) and observed after 24 h of culture with staining by Calcein AM (green; marker for viability) and propidium iodide (red marker for cell death) (n = 3). (b). Laser confocal photomicrographs of ovarian cancer OVCAR-3 cells allowed to transit from moruloid to blastuloid spheroids (transition 2; top represents a cartoon representation of the same). Spheroids were untreated (top row of micrographs) or treated with 250 *μ*M oseltamivir (second from top), 500 *μ*M oseltamivir (third from top), and 2 mM EGTA (bottom row) and observed after 24-72 h of culture with staining by Calcein AM (green; marker for viability) and propidium iodide (red marker for cell death) (n = 3). Scale bar: 50 *μ*m.

**Figure 7:**
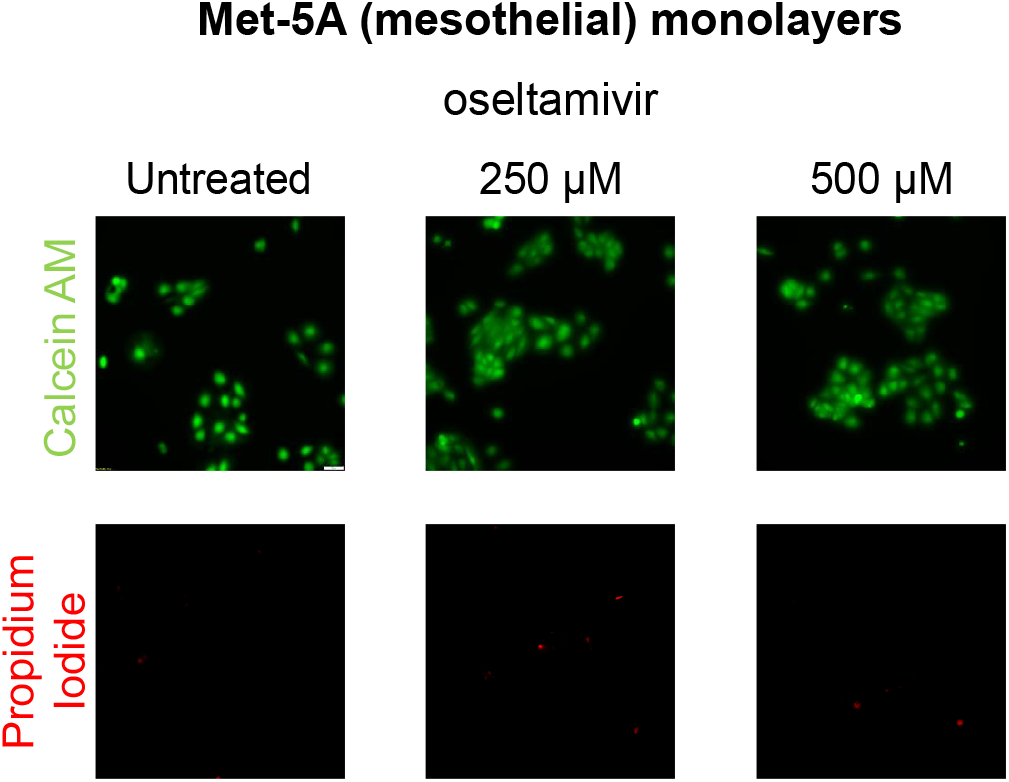
Spheroid disrupting doses of oseltamivir are not cytotoxic to mesothelial monolayers: (a). Laser confocal photomicrographs of the untransformed mesothelial line Met-5A cultivated as monolayers: untreated (left) or treated with 250 *μ*M oseltamivir (middle), 500 *μ*M oseltamivir (right), and observed after 24 h of culture with staining by Calcein AM (green; marker for viability; top row) and propidium iodide (red marker for cell death; bottom row) (n = 3).

## Discussion

Metastatic spheroidal morphologies during ovarian cancer are associated with disease recurrence, drug resistance, and poor overall survival. These spheroidal morphologies synergize with altered cellular metabolism for their energy demands. The molecular profiling of different stages of ovarian cancer uncovers the mechanisms involved in cancer progression. By acquiring protein expression data of different morphological stages of the OVCAR-3 cell line, we identified differentially expressed proteins and observed substantial involvement of enzymatic genes. The results showed high expression of GLS and DBI during transition 1, which previously correlated with poor survival and chemotherapy resistance in ovarian cancer cells [36, 37]. Overexpression of metabolic genes, namely, ILK and MAP2K1 during transition 2, have also been previously associated with tumor metastasis and carboplatin resistance in ovarian tumor cells [38, 39]. Further, pathway analysis of the perturbed proteins revealed the significant involvement of the HIF-1 signaling pathway, glycolysis, and estrogen signaling pathway during both transitions. HIF-1 signaling and metabolic pathways such as glycolysis follow a sequence of signaling cascades to help cancer cells survive and evade the immune response [40]. Essentially metabolic alterations are definitive signatures for cancer progression, and therefore next, we aimed to explore the connection between metabolic reprogramming and morphological stage transition in ovarian cancer.

Metabolic models specific to each morphological stage of OVCAR-3 were built by integrating the corresponding proteomics data into Recon3D. The metabolic pathway enrichment analysis of reactions perturbed during morphological transitions suggested that most of the significantly altered pathways were involved in carbohydrate metabolism, aminoacid/protein metabolism, and lipid metabolism. Both transitions showed a considerable overlap among metabolic perturbations, which essentially included the TCA cycle, fatty acid oxidation & synthesis, and cholesterol metabolism. However, few transition-specific metabolic perturbations were also observed, including alterations in pyruvate metabolism and triacylglycerol synthesis during transition 2. High pyruvate uptake has not only been correlated with invasiveness but also in conjugation with the TCA cycle results in high ATP production, which may serve as a niche for energy demands [41]. In concordance with our findings, high content of triacylglycerols has also been reported in late-stage tumor samples collected from peritoneum [42]. These findings establish the metabolic basis of phenotypic differences in early and late stage transition in OVCAR-3.

We next aimed at bringing down multiple perturbations during disease progression by finding core driver modules. To accomplish this, we devised a new approach in which we clustered perturbed reactions on the basis of their coupled effect on the transition. The top perturbed reaction modules during both the transitions were involved in pathways such as exchange/demand reactions, transport extracellular, transport mitochondrial, and pyrimidine catabolism. During cancer, many mitochondrial transport proteins that link the mitochondrial matrix and cytosol pathways are disrupted [43]. Carbohydrate metabolism pathways such as the TCA cycle, pyruvate metabolism, and pentose phosphate were among the top disease driving modules in both the transitions, underlying an axis of energy requirement during the initial and later stages of ovarian cancer [44, 45, 46]. Transition 1 specific reaction modules were enriched in glycolysis/gluconeogenesis, fatty acid oxidation, and synthesis. In attachment-free environments, ovarian cells shift their metabolic demands on fatty acids to survive. This increased dependency on fat as an energy source causes ovarian cancer cells to resist anoikis (apoptosis due to cell detachment) [47]. The tri-acyl glycerol (TAG) pathway was predominantly involved in the top reaction modules of transition 2. Moreover, the TAG pathway has been known to promote cancer growth by acquiring free fatty acids from tri-acyl glycerides to the cellular pool of fatty acids [48, 49]. During transition 2, arginine and proline metabolism was involved in top modulation strategies and is known to influence collagen synthesis, resulting in increased cancer cell plasticity [50]. Hence, controlling or suppressing perturbations involved in the reactions of the pathways mentioned above can immobilize the progression of the disease towards its more metastatic stage.

Genome-level understanding of metabolic reprogramming by cancer cells may disclose new metabolic targets for drug development or repositioning. Single gene knockout was performed using genes that catalyze metabolic reactions altered during morphological transitions and are not essential for normal cell proliferation. During transition 1, the knockout of genes such as DLST, GALE which catalyze reactions of the pathways involved in carbohydrate metabolism, showed a significant role in reverting the altered metabolic profile. In ovarian cancer, DLST, a component of the alpha-ketoglutarate dehydrogenase complex, plays a vital function in the immuno-metabolic control of immunosuppressive myeloid cells [51]. During transition 2, knocking down SLC transport proteins such as SLC17A4, SLC17A2, SLC25A1, etc., showed a significant role in reverting the metabolic perturbation involved. Over-activation of SLC25A1 has been found in ovarian cancer and inhibiting it improved cancer cell sensitivity to platinum-based chemotherapy [52]. It highlights the fact that cancer cells in 3D (spheroids) culture are more chemo-resistant than cancer cells in 2D (monolayer) culture, which is in accordance with the previous studies [53, 54]. Thus, silencing these drug membrane transporters may help reverse the drug resistance phenotype. The tri-acyl glycerol pathway (involved in lipid metabolism) was prominently involved in top disease driving modules of transition 2, and the knockout of genes, such as GNPAT, GPAM involved in this pathway were able to reverse transition 2. Also, it has been previously observed that GPAM silencing reduced cell migration and tumor xenograft development in ovarian cancer cells [55]. Thus, the TAG pathway plays a significant role in the progression of the disease to its metastatic stage, and the genes involved in this pathway hold considerable potential to become metabolic targets. Single gene knockout of ALDOB, ADH1B, NEU1, and NT5E reduced the metabolic alterations during both transitions. Our results concord with previous studies, which revealed that silencing ADH1B, NEU1, and NT5E can suppress ovarian cancer growth [31, 56, 57].

Drug repurposing opens a feasible space for devising a single/multi-gene targeting strategy in different indications. We evaluated the impact of metabolism-targeting drugs retrieved from the DrugBank database using metabolic models of spheroidal morphologies. The target of disulfiram catalyzes reactions that were perturbed during transition 1, and it has previously been demonstrated to have an antineoplastic effect *in vitro* against ovarian cancer cells [58]. In a previous study, clodronate therapy was found to reduce tumor angiogenesis in an orthotopic ovarian cancer model [59]. This is consistent with our result as clodronate target genes of mitochondrial transport pathway, which was disrupted during both transitions. Canagliflozin, moexipril, oseltamivir, myo-inositol, glyburide, gemcitabine, pentoxifylline, and salicylic acid target proteins that catalyze reactions perturbed during both transitions. Among these drugs, oseltamivir showed larger overlap values and higher pathway perturbation. Oseltamivir is an anti-viral drug indicated for the treatment of influenza A and B virus; it inhibits the neuraminidase enzyme in humans [60]. The metabolic perturbed reactions catalyzed by neuraminidase enzyme were enriched in pathways, namely, keratan sulfate degradation, chondroitin sulfate degradation, N-glycan degradation, sphingolipid metabolism. It was observed that oseltamivir treatment disrupted the single cell to moruloid, and moruloid to blastuloid transitions of ovarian cancer spheroids in a dose dependent fashion, resulting in smaller and dysmorphic spheroids at concentrations of 250 *μ*M. Interestingly, no cytotoxicity was observed either in the cancer cells whose organization was disrupted or monolayers of mesothelia: the untransformed stromal cells of the micro-environment.

The current study aimed to capture the crosstalk between metastatic morphological transitions and cognate metabolic reaction fluxes. The study focused on predicting the transition-specific perturbed metabolic reaction modules responsible for disease progression and identifying regulatory points that could revert the disease.

### Limitations of the study

Though the model predictions show promising results in preliminary *in vitro* experiments, the study needs further confirmation through *in vivo* validation. Since the main focus of the present study is to develop a computational methodology using GSMM, we chose to confirm results through *in vitro* validation. However, the promising results obtained in the *in vitro* study opened new directions for researchers to take the results for further validation and evaluation through *in vitro* and *in vivo* experiments. Further, we chose oseltamivir for *in vitro* validation of our model predictions because of its larger overlap values and higher pathway perturbation. However, the study provides more drugs that restore metabolic perturbations during both transitions. Though they have less potential, per our result, it is still worth checking their effect *in vitro* and *in vivo* for further filtration. Despite these limitations, the current work expands our horizon on ovarian cancer progression and provides a methodological framework for researchers to identify novel targets against cancer progression.

## Highlights

- Quantitative proteomics data produced for distinct morphologies of OVCAR-3 cell line.
- Captured perturbed reaction modules associated with disease progression using GSMM.
- Oseltamivir identified as a potential candidate for impairing cancer progression.
- *In vitro* data shows oseltamivir disrupts morphological transitions of spheroids.

## Declarations

### Ethical approval

The studies described here were carried out in accordance with the guidelines of IISc Bioethics and Safety Committee.

### Availability of data and materials

Supplementary methods, results, tables and figures are provided in pdf file named as Supplementary Information. The datasets generated and analysed during the current study are provided in the link:

https://drive.google.com/drive/folders/1qehnm2s6yP3Bi3gocJlq4y9MGfjMpBsH?usp=sharing

### Conflicts of interest declaration

The authors declare that they have no conflicts of interest.

### Competing Interest

The authors declare that they have no competing interest.

### Funding

The work is supported by SERB (Govt. of India), grant no.: CRG/2019/005477. GA is supported by the THSTI PhD Fellowship. The work of RB is supported by the India Alliance DBT Wellcome Trust Fellowship [IA/I/17/2/503312], the Department of Biotechnology, India [BT/909PR26526/GET/119/92/2017] and SERB [ECR/2015/000280]. RB also acknowledges the financial support of the John Templeton Foundation (#62220). The opinions expressed in this paper are those of the authors and not those of the John Templeton Foundation. JL is supported by Senior Research Fellowship (SRF) from the Ministry of Education, India. MB is supported by the DBT JRF fellowship from the Department of Biotechnology, India.

## Author contributions

SC and RB conceived and designed the study. RB, JL and MB designed the experiments. JL and MB performed the experiments. GA, RB, JL and MB analysed the experimental results. GA designed the model and performed the analysis. GA and SC interpreted the model results. GA wrote the manuscript. SC, RB, JL and MB edited the manuscript. SC coordinated and supervised the work.

## Acknowledgments

Not applicable

## Notes

### Competing Interest Statement

The authors have declared no competing interest.

