## Supplementary Information for "Targeting metabolic fluxes reverts metastatic transitions in ovarian cancer"

### S1 Methods

#### S1.1 Mass spectrometry

In this study, the ovarian cancer cell line OVCAR-3 (obtained from ATCC, Virginia, US) was used. The protocol employed in our previous study [1], was followed in order to maintain and culture the cells in suspension. The 2D cell culture of OVCAR-3 (monolayer) in suspension aggregated into dysmorphic solid clusters, defined as moruloid. These spheroids allowed intercellular movement and penetration by new cells. Moruloid clusters then combined to form bigger clusters that matured further into blastuloid. These spheroids neither combined nor allowed penetration by new cells. Quantitative mass spectrometry was performed on three independent biological replicates of each of the three conditions. The detailed protocol used for mass spectrometric analysis is present in the study [1]. Briefly, samples were reduced with tris(2-carboxyethyl) phosphine (TCEP), alkylated with iodoacetamide, and then digested with Trypsin. Experiments were carried out with EASY-nLC 1,000 system (Thermo Fisher Scientific) paired with a Thermo Fisher-QExactive equipped with a nanoelectrospray ion source. The spectrometric data was collected using a data-dependent top10 approach. The RAW files that were generated were subsequently compared to the Uniprot HUMAN reference proteome database using Proteome Discoverer (v2.2). The false discovery rate for the peptide spectrum match and the protein was set to 0.01 FDR. Statistical analysis was carried out with the help of an in-house R script. The abundance values for each run (containing all biological replicates) were filtered and imputed using a normal distribution.

#### S1.2 Data pre-processing

The high throughput proteomics data contains the expression of 3122 proteins across triplicates of the distinct phenotypic morphologies monolayer, moruloid spheroids, and blastuloid spheroids. We first removed proteins absent in all the nine samples of the data. Next, 2720 proteins expressing in 50% of the replicates in at least one morphological stage were filtered out. For missing value imputation, proteins expressed in at least 50% of the replicates, the missing protein expression was replaced by the mean value of the protein expression in the other two replicates. Otherwise, it was replaced by the minimum of the data. Next, the protein expression of replicates in each morphological stage was merged by taking the mean of the protein expression values of all its replicates. Gene names corresponding to the proteins present in the data were intersected with genes present in Recon3D, and 604 metabolic genes were filtered out. Finally, the data was $\log_{10}$ normalized and the metabolic genes data was further used for building the models.

#### S1.3 Steps for building context-specific metabolic models

##### S1.3.1 Fixing reaction flux bounds

We constrained the carbon-source exchanges to a maximum uptake of 10 mmol $gDW^{-1}h^{-1}$ [2]. Other reactions were set to the original flux bounds of the Recon3D model.

##### S1.3.2 Mapping of expression data to reactions

In order to determine the expression data associated with each metabolic reaction present in Recon3D, we used the GPR rules present in the model. GPR rules describe the association between the reaction and the gene product catalyzing it. A function named 'GPRparser' of COBRA Toolbox [3] was used to map the GPR rules to a specific format that can further be used. Then, the cell-matrix containing the parsed GPR rule was used to integrate the expression data and associate it to each reaction in the model using the 'mapExpressionToReactions' of the COBRA Toolbox. This resulted in a reaction expression array, which had an expression for each reaction.

Finally, using the following three algorithms and gene expression thresholds (mentioned in each algorithm), we built the context-specific metabolic models for each morphological stage.

###### S1.3.3 Integrative Metabolic Analysis Tool (iMAT)

The iMAT implementation [4] in COBRA Toolbox was used to extract context-specific metabolic models using expression data. The algorithm takes the following inputs:

1. genome-scale metabolic model (Recon3D),
2. expression data of reactions (see section S1.3.2),
3. lower threshold (i.e., reactions with expression value below this value were considered to be non-expressed),
4. upper threshold (i.e., reactions with expression value above this value were considered to be expressed)

The algorithm aims to include the high expression reactions and remove the low expression reactions while building the model. The two threshold values used in our study for this algorithm are:

Threshold 1 (Th1): lower expression threshold: 2.4861 (minimum value of reaction's expression array), and upper expression threshold: minimum value of top 10% of reaction's expression array
Threshold 2 (Th2): lower expression threshold: mean(exp) − std(exp), and upper expression threshold: mean(exp) + std(exp)
where, 'exp' is the expression array of reactions in the model, and 'std' is the standard deviation

###### S1.3.4 FASTCORE

FASTCORE is an algorithm to build context-specific models with the advantage of fast computation and compactness of the output model [5]. The algorithm aims to determine a set of core reactions using gene expression threshold value that are active in the output model. The algorithm also finds a minimum number of reactions to support the set of core reactions in order to make the final model consistent. The reactions with no expression data were not considered core reactions. The two threshold values used in our study for this algorithm are:

Threshold 1 (Th1): minimum value of top 10% of reaction's expression array
Threshold 2 (Th2): 2.4861 (minimum value of reaction's expression array)

##### S1.3.5 Integrative Network Inference for Tissues (INIT)

INIT is a model extraction algorithm [6], that includes and removes reactions based on the weights allotted to them, represented mathematically as:

$$\text{weight} = 5 \times \log_2 \left( \frac{\text{Expression level}}{\text{Threshold}} \right)$$

where, Expression level is the reaction's expression evaluated using GPR rules.
Reactions with positive weights were considered to have high expression, whereas reactions with negative weights were considered as less expressed. A weight of $-2$ was given to reactions with no expression data. The algorithm ascertains the presence of highly expressed reactions by maximizing the sum of the weights of reactions. The two threshold values used in our study for this algorithm are:

Threshold 1 (Th1): minimum value of top 10% of reaction's expression array
Threshold 2 (Th2): 2.4861 (minimum value of reaction's expression array)

The complete workflow from constructing context-specific metabolic models to evaluating the performance of each model is shown in Supplementary Fig. S1.

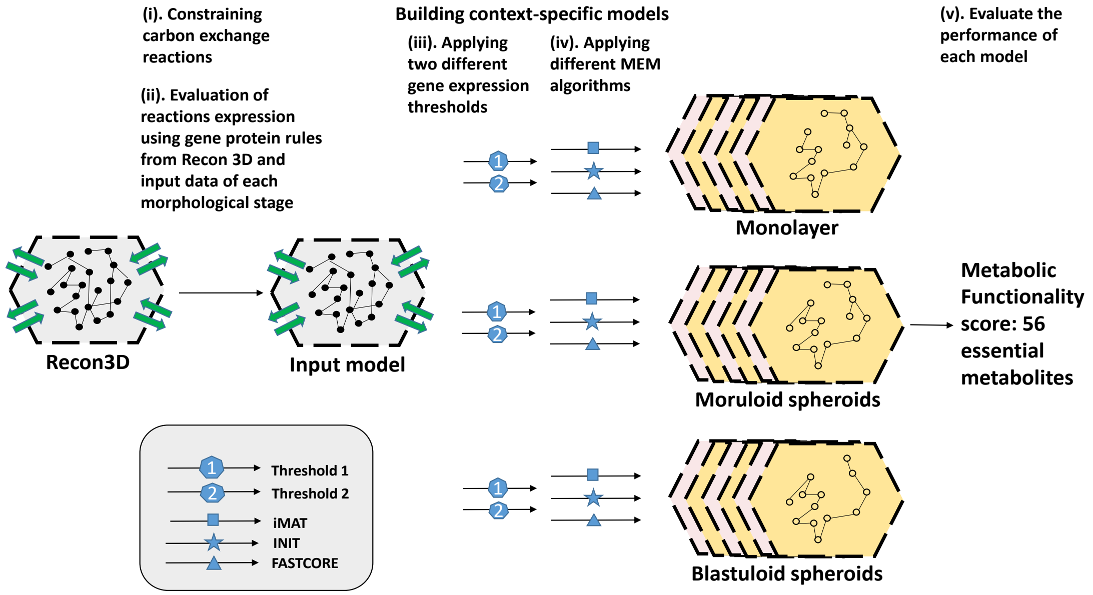


Figure S1: **Model construction workflow:** The figure describes the workflow for the construction of context-specific metabolic models by applying different MEM algorithms, gene expression thresholds and evaluating the functionality of each model.

#### S1.4 Evaluating steady-state flux profiles

By using the constraint-based modeling (CBM) approach as suggested by Tomer et al. [7], we allotted possible flux distribution to all the reactions in the metabolic model in a way that satisfies the stoichiometric and thermodynamic constraints embedded in the model. Flux sampling produces a sequence of feasible solutions (called a chain) that satisfy all the network constraints while analyzing the entire solution space. To account for these solutions, we used a CBM sampling technique, GP sampler, which samples an arbitrary linearly constrained space with the help of a fixed number of points (default = 2 × number of reactions in the model). The space is defined as:

$$Ax = b$$

$$lb \leq x \leq ub$$

The obtained solution space from the linear constrained problem is represented by:

$$E = \{(f_{1,j}, f_{2,j}, ..., f_{n,j}) : f_{i,j} \in \mathbb{R}, i = 1, 2, ..., n\}$$

where, $n$= no. of reactions, $j$= fixed no. of points

Finally, out of the sampled solution space, the solution having a maximum correlation with the reaction's expression array (see section S1.3.2) was regarded as the optimal flux solution, which is denoted by:

$$F = \{(f_1, f_2, ..., f_n), \quad f_i \in E, \ \max(\mathrm{corr}((f_i), \mathrm{exp}))\}$$

where, $i = 1, 2, ..., n$, $n$= number of reactions, exp= reaction's expression array, corr(($f_i$), exp)= correlation between flux values and expression array of metabolic reactions in the queried model.

### S2 Results

#### S2.1 OVCAR-3 proteomics data reveal perturbations associated with morphological transitions

High throughput proteomics was performed in this study using the mass spectrometry data-dependent top10 approach. The data contains the expression of 3122 proteins across the three biological replicate samples of each morphological stage of the OVCAR-3 cell line (see Fig. S2a), namely, monolayer, moruloid (24-hours spheroids), and blastuloid (1-week spheroids) (see Methods: S1.1). The proteomics data relating to both the monolayer-moruloid transition and the moruloid-blastuloid transition is provided in Data File 1. A part of this proteomics data related to moruloid-blastuloid transition is also provided in a previous study [1]. The proteomics data was first pre-processed in order to account for missing data and variability (see Methods: S1.2). The protein expressions showed varying perturbation levels during the initial and later stages of the morphological transition of the OVCAR-3 cell line. Agglomerative hierarchical clustering revealed grouping among the replicates of each morphological stage, trailed by clustering in samples of moruloid and blastuloid spheroids (see Fig. S2b).

Differentially expressed proteins between progressive morphological stages of OVCAR-3 were identified by applying a cutoff of 1 on $\log_2$ fold-change of protein expression values (see Fig. S2c). Among the top perturbed genes, GNPNAT1, CTBP1, GMPR, and SLC25A6 were found to be commonly perturbed during both transitions. During transition 1, genes GLS, DBI, and STK4 were among the top perturbed metabolic genes. Moreover, during transition 2, metabolic genes such as ILK, MAP2K1, and AHCYL1 were perturbed. Using Enrichr [8], pathway enrichment analysis of differentially expressed proteins was done for both the transitions (see Data File 2). Metabolic reprogramming is the major hallmark of cancer; however, its role in different morphological stages of OVCAR-3 might help us to understand the tumor progression in ovarian cancer.

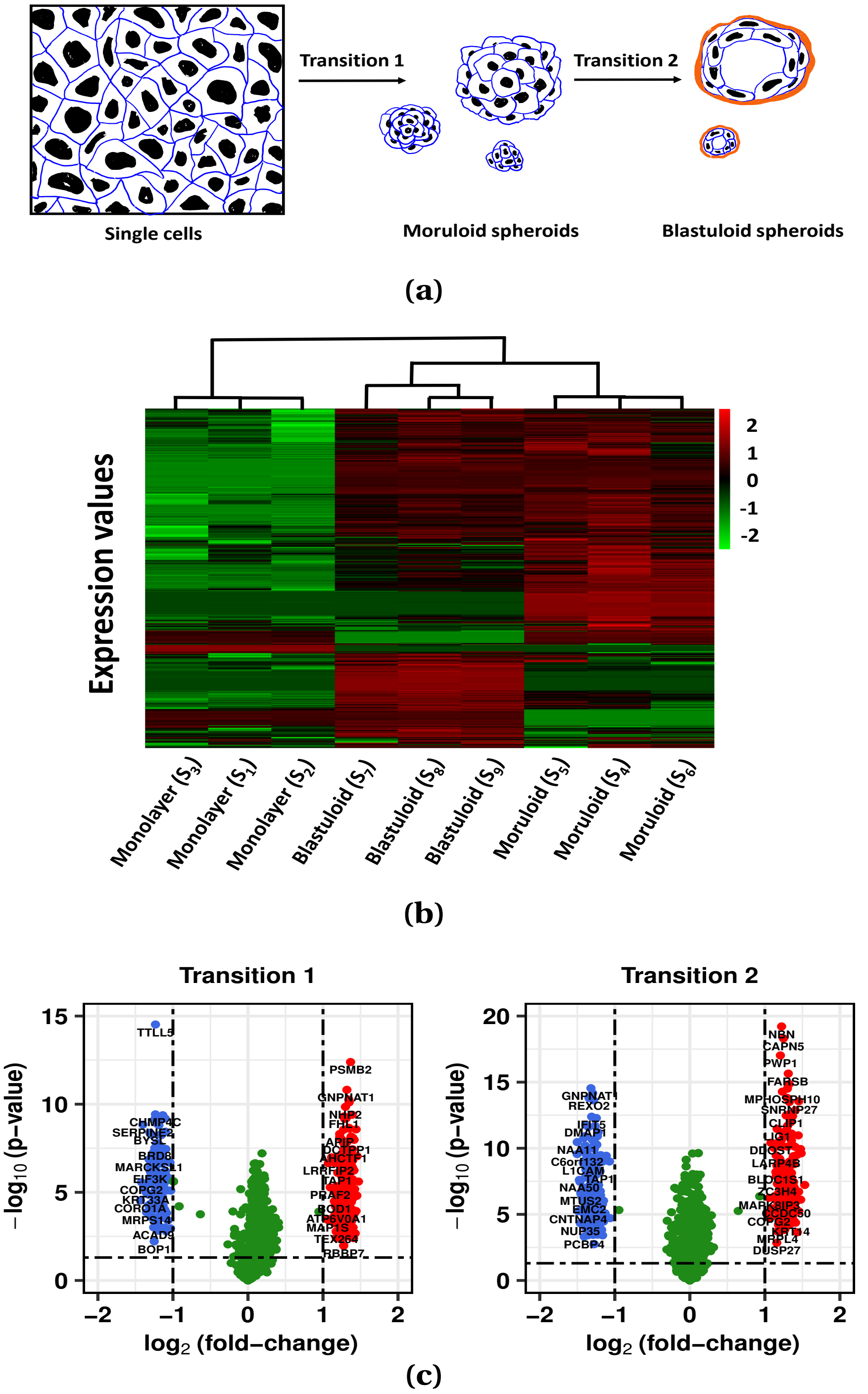


Figure S2: **Proteomic changes during morphological transitions in OVCAR-3 ovarian cancer cells:** (a). Cartoon depiction of the single cell-to-moruloid and moruloid-to-blastuloid transitions. (b). Heat map showing standardized protein expressions across all 9 (3x3) samples. The color bar shows the standardized protein expression value, and the dendrogram represents hierarchical clustering among the samples of morphological stages of OVCAR-3. (c). Volcano plots showing the distribution of differentially expressed proteins for both the morphological stage transitions. The X-axis represents $\log_2$ (fold-change) of expression values between the profiles, and the Y-axis represents -$\log_{10}$ (p-value) obtained from a t-test of differences between the samples. The horizontal dashed line represents p-value $= 0.05$, whereas vertical dashed lines partition the plot for $|\log_2$ (fold-change)$| \geq 1$. Blue and red dots represent the down-regulated and the up-regulated proteins, respectively, whereas green dots represent not differentially expressed proteins.

#### S2.2 Metabolic profiling of the iMAT-Th1 models translates morphological transitions into metabolic perturbations

The iMAT-Th1 models had a high metabolic functionality score and were less sensitive to the threshold selection, therefore, they were considered for further investigation. Constraint-based modeling (CBM) flux sampling technique was used to evaluate the steady-state flux values, and the set of flux values with the highest correlation with the input expression data was considered the optimal solution (see Methods: S1.4). A 2 fold cut-off was applied on flux fold-change values of reactions present in the models of both stages to determine reactions perturbed during each transition. It was found that a total of 1226 and 1252 reactions were perturbed in transition 1 and transition 2, respectively (see Fig. S3a).The flux value of 675 reactions commonly perturbed during both morphological stage transitions has been represented through the heat-map (see Fig. S3b). The clustering among moruloid and blastuloid profiles was retained after the integration of proteomics data into the metabolic model (see Fig. S3b). The metabolic pathway enrichment analysis of perturbed reactions was done using the Recon3D database. The significance of overlap was evaluated using the hypergeometric test. The enriched pathways were further categorized based on their participation in various metabolic processes. Lipid, carbohydrate, and protein/amino-acid metabolism contributed a significant share in the progression of the disease (see Fig. S3c). Several pathways were impaired during both transitions, including glycolysis, TCA, fatty acid synthesis & oxidation, cholesterol metabolism, and a few transport mechanisms. Among nucleotide metabolism-associated pathways, pyrimidine synthesis and purine catabolism were only disrupted during transition 1. Pathways involved in amino acid metabolism such as alanine & aspartate and glycine, serine, alanine, and threonine metabolism were significantly perturbed, specifically in transition 1. On the other hand, for transition 2, specific perturbation in methionine & cysteine, lysine metabolism was observed. During transition 2, alterations in pyruvate metabolism, which is a part of carbohydrate metabolism, were also discovered. Moreover, a few pathways involved in lipid metabolism, namely, tri-acylglycerol synthesis and steroid metabolism, were also significantly altered during transition 2.

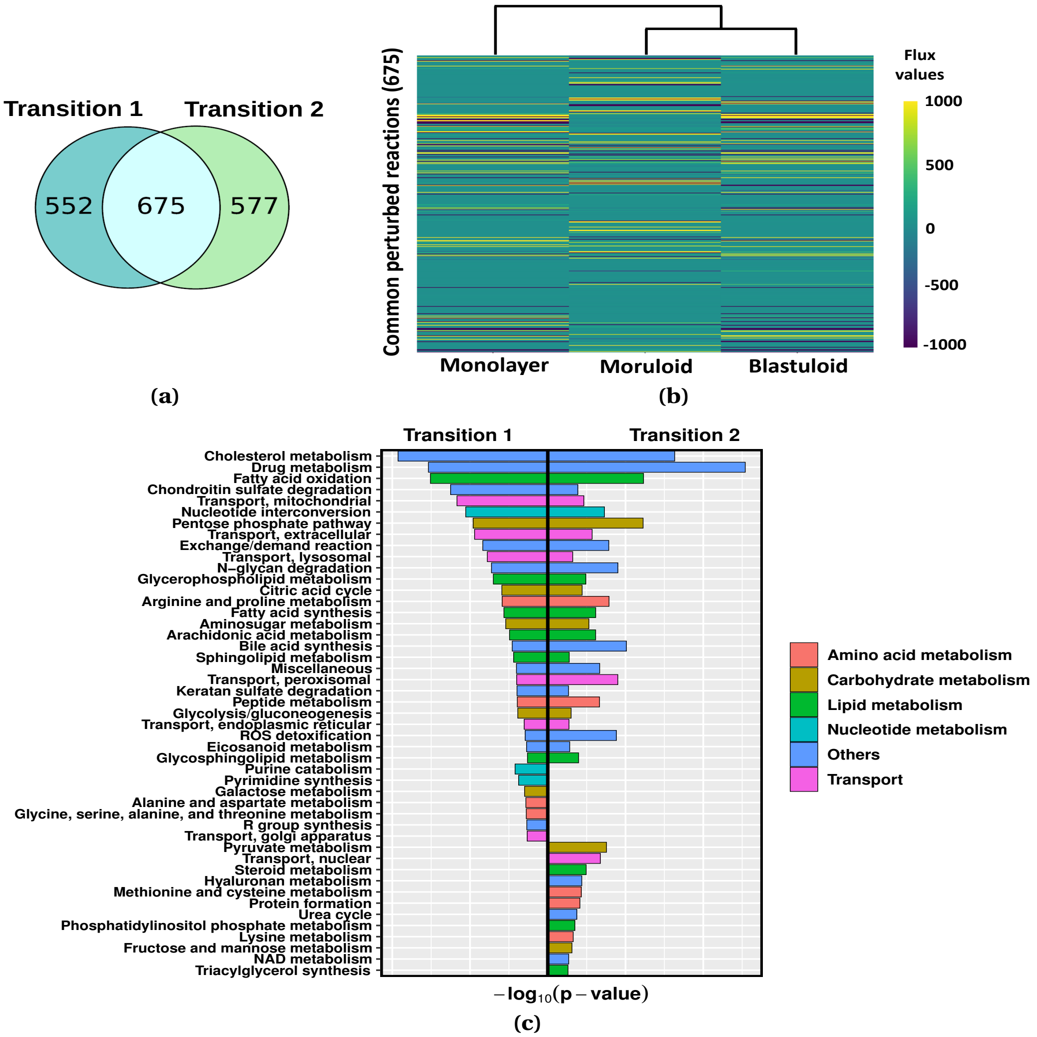


Figure S3: **Metabolic perturbations in steady-state flux profile of iMAT-Th1 models:** (a). Venn diagram shows the number of common and specific reactions perturbed during each transition. (b). Heat map represents the steady-state flux values of the reactions commonly perturbed (675) during both the transitions and the hierarchical clustering among the metabolic profiles of the morphological stages. (c). The figure shows the pathways significantly perturbed (p-value $< 0.05$) during both transitions. The color coding represents the presence of pathways involved in different metabolic processes.

#### S3 Tables

Table S1: **Targetibility score of top 20 modulation strategies of transition 1**

| Strategies | Targetibility Score |
|---|---|
| **A1** | 84 |
| **A2** | 82 |
| **A3** | 61 |
| **A4** | 79 |
| **A5** | 68 |
| **A6** | 79 |
| **A7** | 65 |
| **A8** | 83 |
| **A9** | 68 |
| **A10** | 83 |
| **A11** | 71 |
| **A12** | 80 |
| **A13** | 69 |
| **A14** | 64 |
| **A15** | 73 |
| **A16** | 73 |
| **A17** | 73 |
| **A18** | 68 |
| **A19** | 76 |
| **A20** | 76 |

Table S2: **Targetibility score of top 20 modulation strategies of transition 2**

| Strategies | Targetibility Score |
|---|---|
| **B1** | 79 |
| **B2** | 77 |
| **B3** | 50 |
| **B4** | 63 |
| **B5** | 86 |
| **B6** | 77 |
| **B7** | 73 |
| **B8** | 71 |
| **B9** | 50 |
| **B10** | 75 |
| **B11** | 75 |
| **B12** | 90 |
| **B13** | 71 |
| **B14** | 67 |
| **B15** | 64 |
| **B16** | 74 |
| **B17** | 59 |
| **B18** | 65 |
| **B19** | 83 |
| **B20** | 91 |

Table S3: **In-silico validation using GDSE sensitivity score (transition 1)**

| Drugs | Drug sensitivity score | Correlation after drug addition |
|---|---|---|
| **Afatinib** | -0.019774 | 0.1963 |
| **Bosutinib** | 0.98623 | 0.0309 |
| **Cetuximab** | 0.36838 | 0.0276 |
| **Crizotinib** | 1.07 | 0.0276 |
| **Cytarabine** | 0.70312 | 0.0841 |
| **Dabrafenib** | 0.6468 | 0.0276 |
| **Dasatinib** | -0.50914 | 0.1963 |
| **Gemcitabine** | 0.92802 | 0.2155 |
| **Ibrutinib** | 0.34828 | 0.0276 |
| **Lapatinib** | 1.0507 | 0.1963 |
| **Leflunomide** | 0.26131 | 0.2508 |
| **Pazopanib** | -0.29808 | 0.2101 |
| **Pemetrexed** | 0.44992 | 0.0903 |
| **Ponatinib** | 0.42051 | 0.0276 |
| **Ruxolitinib** | 0.37862 | 0.0276 |
| **Sorafenib** | -0.17518 | 0.1931 |

Table S4: **In-silico validation using GDSE sensitivity score (transition 2)**

| Drugs | Drug sensitivity score | Correlation after drug addition |
|---|---|---|
| **Afatinib** | -0.019774 | 0.2051 |
| **Bosutinib** | 0.98623 | 0.0389 |
| **Cetuximab** | 0.36838 | 0.1887 |
| **Crizotinib** | 1.07 | 0.0389 |
| **Cytarabine** | 0.70312 | 0.0352 |
| **Dabrafenib** | 0.6468 | 0.1887 |
| **Dasatinib** | -0.50914 | 0.2033 |
| **Gemcitabine** | 0.92802 | 0.1851 |
| **Ibrutinib** | 0.34828 | 0.1887 |
| **Lapatinib** | 1.0507 | 0.2051 |
| **Leflunomide** | 0.26131 | 0.1372 |
| **Pazopanib** | -0.29808 | 0.1993 |
| **Pemetrexed** | 0.44992 | 0.1537 |
| **Ponatinib** | 0.42051 | 0.1887 |
| **Ruxolitinib** | 0.37862 | 0.1887 |
| **Sorafenib** | -0.17518 | 0.2088 |

#### S4 Figure

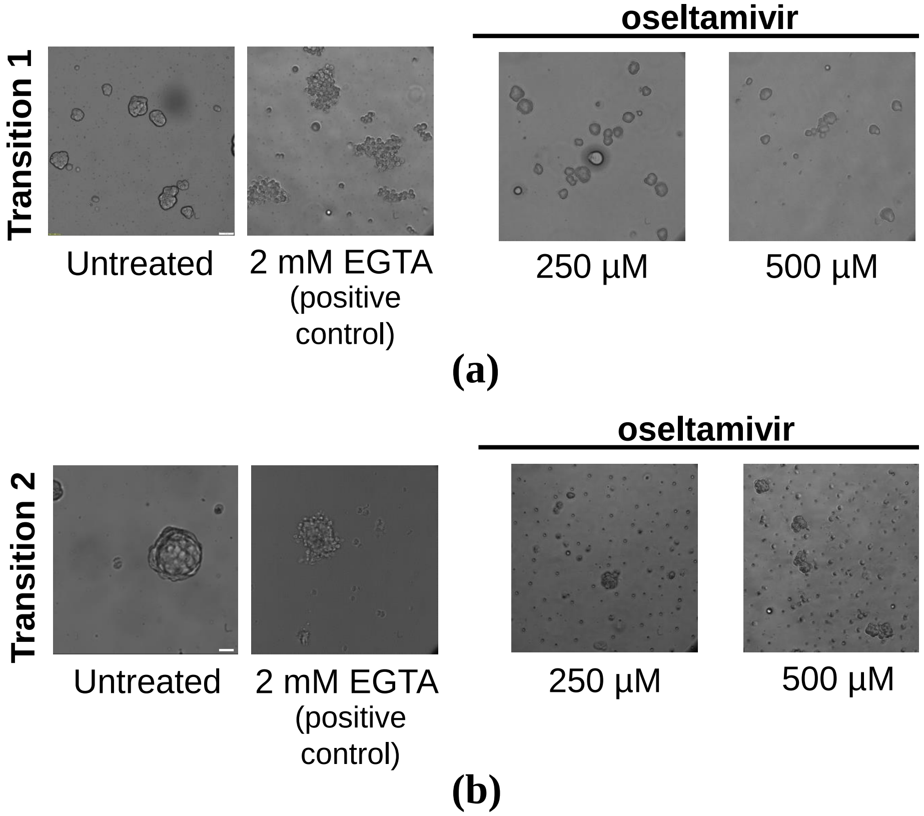


Figure S4: **Oseltamivir treatment disrupts single cell-to-moruloid and moruloid-to-blastuloid transitions of ovarian cancer spheroids:** (a). Phase contrast photomicrographs of ovarian cancer OVCAR-3 cells allowed to transit from single cells to moruloid spheroids (transition 1). Cells were untreated or treated with 250 $\mu$M and 500 $\mu$M oseltamivir and 2 mM EGTA and observed after 24 h of culture (n = 3). (b). Phase contrast photomicrographs of ovarian cancer OVCAR-3 cells allowed to transit from moruloid to blastuloid spheroids (transition 2). Spheroids were untreated or treated with 250 $\mu$M and 500 $\mu$M oseltamivir and 2 mM EGTA and observed after 24-72 h of culture (n = 3). Scale bar: 50 $\mu$m.
